# KLRG1 marks tumor-infiltrating CD4 T cell subsets associated with tumor progression and immunotherapy response

**DOI:** 10.1101/2023.01.01.522340

**Authors:** Casey R. Ager, Mingxuan Zhang, Matthew Chaimowitz, Shruti Bansal, Aleksandar Obradovic, Meri Rogava, Johannes C. Melms, Patrick McCann, Catherine Spina, Charles G. Drake, Matthew C. Dallos, Benjamin Izar

## Abstract

Current methods for biomarker discovery and target identification in immuno-oncology rely on static snapshots of tumor immunity. To thoroughly characterize the temporal nature of antitumor immune responses, we developed a 34-parameter spectral flow cytometry panel and performed high-throughput analyses in critical contexts. We leveraged two distinct preclinical models that recapitulate cancer immunoediting (NPK-C1) and immune checkpoint blockade (ICB) response (MC38), respectively, and profiled multiple relevant tissues at and around key inflection points of immune surveillance and escape and/or ICB response. Machine learning-driven data analysis revealed a pattern of KLRG1 expression that uniquely identified intratumoral effector CD4 T cell populations that constitutively associate with tumor burden across tumor models, and are lost in tumors undergoing regression in response to ICB. Similarly, a Helios^-^KLRG1 ^+^ subset of tumor-infiltrating regulatory T cells (Tregs) was associated with tumor progression from immune equilibrium to escape, and were also lost in tumors responding to ICB. Validation studies confirmed KLRG1 signatures in human tumorinfiltrating CD4 T cells associate with disease progression in renal cancer. These findings nominate KLRG1^+^ CD4 T cell populations as subsets for further investigation in cancer immunity and demonstrate the utility of longitudinal spectral flow profiling as an engine of dynamic biomarker and/or target discovery.

## Introduction

Immunotherapy is now a pillar of cancer treatment. However, responses to most immunotherapeutic agents remain rare, restricted to a limited number of tumor types, and difficult to predict(1). Improving response rates and developing biomarkers predictive of response are central goals of the tumor immunology field, but this remains challenging. A multitude of complex multi-cellular interactions may govern the outcome of an antitumor immune response in any given patient. Therefore, it is common that single biomarkers – or therapeutic modulation of singular pathways – tend to have utility in some, but not most cancer patients.

For example, there are three immune-related biomarkers currently in widespread clinical use; (i) tumor mutational burden (TMB), as measured directly or inferred via the presence of disabling mutations in DNA repair machinery(2–4), (ii) PD-L1 expression by immunohistochemistry (IHC)(5), and (iii) pre-existing immunity as measured by the presence of CD8 T cells within and surrounding tumors, termed the Immunoscore(6). While each is significantly associated with response to immunotherapy in certain contexts, the sensitivity and specificity of these predictors remain sub-optimal(7). These unimodal approaches likely fail to fully capture the complex mechanisms underlying antitumor immunity, including the dynamic nature of such responses occurring across tissues and over time. As such, application of highly multiplexed single cell immune profiling approaches to biomarker detection and target discovery in a cross-tissue, longitudinal manner may provide nuanced immune phenotypes with greater predictive and translational utility.

To this end, we developed a 34-parameter spectral flow cytometry panel and high dimensional data analysis pipeline to interrogate protein-level immune phenotypes associated with different phases of the cancer immunoediting cycle, including immune escape and tumor outgrowth, and immune checkpoint blockade (ICB) response across preclinical models. We profiled the NPK-C1 and ICB-treated MC38 models, analyzing multiple tissues (tumor, draining versus non-draining lymph nodes, and blood) over and around key inflection points in tumor progression. This facilitated deep characterization of dynamics underlying natural and ICB-induced tumor regression, transition to immune equilibrium, and subsequent transition from equilibrium to uncontrolled tumor escape. Validation studies in human single cell datasets of renal cancer confirmed the translational relevance of these findings.

## Materials and Methods

### Mice

Male and female C57BL/6J mice (5-6 weeks old) were purchased from the Jackson Laboratory (Bar Harbor, ME). Mice were 6-8 weeks old at time of use. All animals were housed in strict accordance with NIH and American Association of Laboratory Animal Care regulations. All experiments and procedures for this study were approved by the Columbia University Medical Center Institutional Animal Care and Use Committee (IACUC).

### Cell Lines

The NPK-C1 cell line (originally LM7304) was provided by Dr. Cory Abate-Shen at Columbia University. See prior references for further detail(8,9). NPK-C1 cells were maintained in R10, consisting of RPMI medium (Corning; Corning, NY) supplemented with 10% FBS (HyClone; Logan, UT), 100 U/mL penicillin, and 100 mg/mL streptomycin (Gibco; Gaithersburg, MD). MC38 colon carcinoma cells were purchased from Kerafast and cultured in D10 consisting of DMEM (Corning; Corning, NY) supplemented with 10% FBS, 100 U/mL penicillin, and 100 mg/mL streptomycin (Gibco; Gaithersburg, MD).

### Tumor Challenge and Therapy Injections

NPK-C1 cells at 70-90% confluence or MC38 cells at 50-75% confluence were harvested with 0.05% trypsin (Gibco, Gaithersburg, MD), washed with PBS, counted, and resuspended at 10×10^6^ cells/mL in ice cold PBS. On day 0, 6-8 week old mice were implanted on the right flank with 1×10^6^ NPK-C1 or MC38 cells. Tumor measurements were recorded in X, Y, and Z dimensions (largest diameter, smallest diameter, and largest height, respectively) every 2-3 days by digital caliper and tumor volume was calculated by multiplying X*Y*Z. Anti-PD-1 (*RMP1-14* IgG2a,κ) or Rat IgG2a isotype antibody (Bio X Cell; Lebanon, NH) was diluted in sterile PBS and administered by intraperitoneal (IP) injection at 200 μg/mL per mouse on days 3, 6, 9, and 12 post-MC38 implantation.

### Tissue Harvesting

Following mouse euthanasia, ~200 μl blood was isolated via cardiac puncture using a 0.2% heparin (StemCell Technologies; Vancouver, BC) coated syringe needle and placed on ice in tubes containing 10 μl 0.5M EDTA (Corning; Corning, NY). Tumor-draining and non-draining inguinal lymph nodes were dissected and placed in 48-well plates containing 150 μl R10 media on ice. Tumors were harvested, massed, and up to 50 mg tumor was diced and placed in X-Vivo 15 media (Lonza; Basel, Switzerland) in 5mL Eppendorf tubes on ice. DNase (40 μl of 20 mg/mL solution; Roche; Basel, Switzerland) and Collagenase D (125 μl of 40 mg/mL solution; Roche; Basel, Switzerland) were added to tumor samples prior to incubation on a shaker at 37° C for 30 minutes. Digest reactions were quenched by vortexing (30 seconds) and adding 5mL R10 media. Tumor digest were filtered through 70 μm filters (Miltenyi; Bergisch Gladbach, GE) and pelleted. Blood samples were RBC lysed with two successive 2 minute incubations in 2mL ACK buffer (Quality Biological; Gaithersburg, MD) and quenching in R10 prior to pelleting. Lymph nodes were physically disaggregated with a 1mL syringe plunger in the 48-well plate, then suspensions were filtered through 40 um filters. All samples were placed in U-bottom 96-well plates for staining.

### Flow Cytometry

Samples were washed with PBS. Dead cells were stained by resuspension in 100 μl PBS + Live/Dead Fixable Blue dye (1:500; Invitrogen; Waltham, MA). This and all further staining or fixation steps were performed for 30 minutes at room temperature on a plate shaker, protected from light. Samples were washed 2x with PBS, then were resuspended in FACS (PBS + 3% FBS + 1mM EDTA + 10mM HEPES) supplemented with TruStain FcX (1:50; BioLegend; San Diego, CA) and TruStain Monocyte Blocker (1:20; BioLegend; San Diego, CA) and placed on ice. Surface antibodies were prepared at optimal dilutions (see Supplementary Table S1) in FACS supplemented with Brilliant Stain Plus buffer (BD; Franklin Lakes, NJ), then were added to samples in blocking solution. After staining, samples were washed 2x with FACS buffer and fixed in 100 μl of FoxP3 Fixation/Permeabilization Kit buffer (eBioscience; San Diego, CA). Samples were washed twice with 1X Permeabilization buffer (1X PW) then were stained in 1X PW plus intracellular antibodies. Samples were then washed twice in 1X PW then fixed in 100 μl of FluoroFix buffer (BioLegend: San Diego, CA), washed twice with FACS, then resuspended in 200 μl FACS and sealed in the 96 well plate for acquisition the following day on a Cytek Aurora 5-laser cytometer. Simultaneously stained splenocyte samples were utilized for single stain controls.

### Analysis and Statistics

Longitudinal profiling experiments were conducted once each for MC38 and NPK-C1 models. A total of n = 10-25 mice per time point were evaluated. Flow cytometry data was analyzed in FlowJo v10.8.1 (BD; Franklin Lakes, NJ) and high dimensional plugin algorithms UMAP and FlowSOM were downloaded from FlowJo Exchange. Up to 30k live CD45+ tumor-infiltrating cells and up to 15k live CD45+ blood or LN-derived cells per sample were concatenated for high dimensional analysis. All dimensionality reduction and clustering were performed in FlowJo. Statistical analyses were performed in GraphPad Prism v9.3.1. Pearson Correlation was used to generate correlation coefficients, and unpaired Student’s T tests with Welch’s correction were used to compare cluster frequencies between groups. Single cell analytical methods are previously described(10,11). The KLRG1 regulon was extracted from the consensus metaVIPER KLRG1 regulon from Obradovic, *et al(11*). To compute the KLRG1 regulon signature, we fit a random forest regression model with regulons genes as features and KLRG1 as target, then computed Gini importance scores of regulon genes to determine their impact on KLRG1 expression. The KLRG1 regulon score of each cell wass given by the weighted sum of expression measures of all regulon genes in the cell with importance scores as weights.

## Results

### Longitudinal profiling captures generalizable features of failed versus successful antitumor immunity

We generated a 34-parameter spectral flow cytometry panel (see list of antibodies used in Supplementary Table S1 and additional clustering in Figure S1) to profile productive vs non-productive immunity in the NPK-C1 model, known to undergo spontaneous immunoediting *in vivo(8*), and the well-characterized MC38 model treated with curative doses of anti-PD-1 immunotherapy (Figure 1A). Specifically, we profiled days 7, 10, and 13 post-implantation of NPK-C1 and MC38, and additional days 20 and 24 of NPK-C1 (representing the equilibrium to escape transition). We utilized a rigorous high dimensional analysis pipeline improved upon a previous study(8) with minimally supervised UMAP (Uniform Manifold Approximation Projection) dimensionality reduction and FlowSOM clustering algorithms to establish cellular identities in high dimensional space (see Figure S1A-E for analysis workflow).

**Figure 1.**
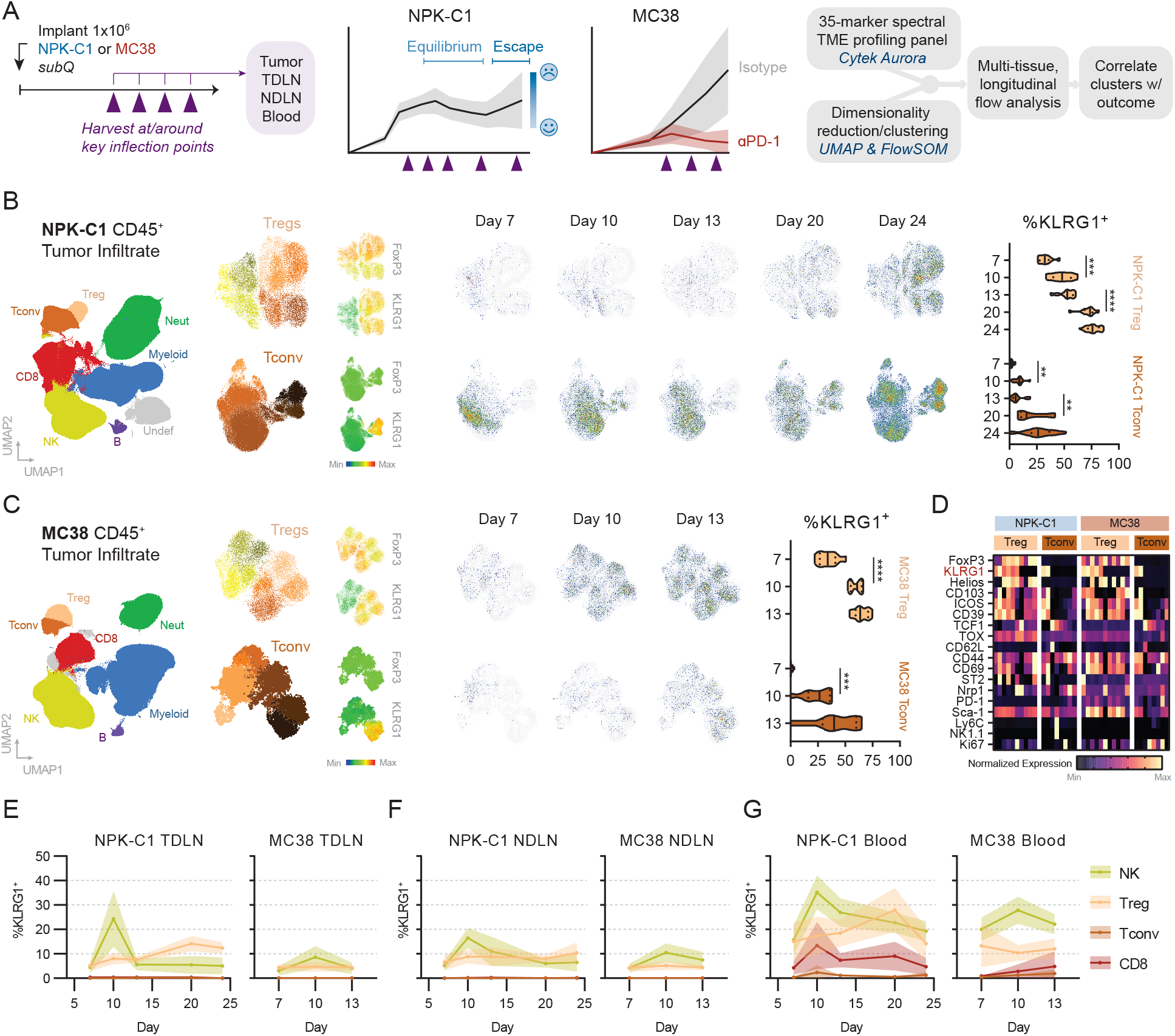
Expression of KLRG1 on tumor-infiltrating CD4 T cells over time in MC38 and NPK-C1. (A) Schematic summary of study design, models used, and analysis pipeline. (B) Dimensionality reduction (UMAP) and clustering (FlowSOM) of NPK-C1 (B) and MC38 (C) infiltrating immune cells, focusing on CD4+ Treg and Tconv populations. Cell-type subclustering, expression of FoxP3 and KLRG1 by heat map coloring, UMAP time courses, and frequencies of KLRG1 expression in each cell type. (D) Expression of phenotypic markers on all CD4 T cell sub-clusters across NPK-C1 and MC38 markers. Color scale is normalized to min/max of the cell type or all immune cells, as appropriate. Quantitation of KLRG1 expression in other immune cell types in TDLN (E), NDLNs (F), and blood (G). Data were generated from one experiment of n=10-25 mice per/condition and time point. Statistical significance is denoted by * (p<0.05), ** (p<0.01), *** (p<0.001), **** (p<0.0001).

### KLRG1 accumulates selectively on tumor-infiltrating CD4 T cell subsets

We observed diverse phenotypic heterogeneity in tumor-infiltrating Tregs infiltrating both NPK-C1 and MC38 tumors (Figure 1B, S1B-C). In both models, we observed a dynamic shift in these phenotypes as tumors progressed. At late time points, up to 80% of Tregs existed in clusters marked by expression of KLRG1 (Fig 1B-C). We observed a similar phenotypic trajectory in the FoxP3^-^ Tconv compartment, as KLRG1 expression defined a distinct Tconv cluster accounting for up to 50-70% of tumor Tconv at late time points in NPK-C1 and MC38, respectively. Phenotypic analysis of KLRG1 ^+^ versus KLRG1^-^ clusters from both tumor models indicated a consistent co-expression pattern of KLRG1 with high levels of CD39 and ICOS on both tumor-infiltrating Tregs and Tconv (Fig 1D). In contrast, ST2/IL-33R, CD69, and CD103 were present on subsets of both KLRG1 ^+^ and KLRG1^-^ Tregs and Tconv, suggesting KLRG1 marks CD4 cell states independent of tissue residence programming or IL-33 signaling, as has been reported(12).

To determine whether KLRG1 is selectively induced on tumor-infiltrating CD4 T cells during tumor progression, we measured expression of KLRG1 on other immune subsets as well as CD4 T cells in tumor-draining (TDLN) and non-draining lymph nodes (NDLN) and blood (Fig 1E). KLRG1 was expressed by 5-10% of Tregs in both TDLNs and NDLNs, and throughout tumor progression a moderate trend towards an increase in KLRG1 ^+^ Tregs was observed in the TDLN, but not the NDLN. NK cells were the only other subset to express KLRG1 in LNs, however this expression did not increase with tumor progression. In the blood, KLRG1 was expressed on subsets of CD8s, Tregs, Tconv, and NK cells, however the frequency of cells expressing KLRG1 in each compartment did not correlate with time of tumor progression. Together, these data reveal a tumor model-independent and TME-restricted phenomenon in which CD4 T cell subsets uniquely gain increasing levels of KLRG1 expression throughout tumor progression.

### Lack of tumor KLRG1^+^ Tconv associates with enhanced tumor control

Based on these data, we next asked whether KLRG1 expression on tumor CD4 T cells has utility as a biomarker of tumor burden or ICB response. We first generated Pearson correlation coefficients for all predominant immune phenotypes in the TME across time points and both treatment-naïve models (Figure 2A). We manually curated these phenotypes into 22 immune metaclusters by grouping FlowSOM-derived subpopulations sharing a dominant sub-phenotypic marker. While the majority of clusters varied in association with tumor burden over time and across models, KLRG1 ^+^ Tconv were strikingly consistent in their correlation with increased tumor burden at every time point in both NPK-C1 and MC38 (Figure 2A-B). On day 24, NPK-C1 tumors either begin progressing to immune escape or stay under immune-regulated equilibrium, allowing assessment of cell types implicated in the poorly understood but clinically-relevant equilibrium-to-escape transition (Figure 2C)(8). Of note, KLRG1 ^+^ Tconv cells were highly enriched in NPK-C1 tumors escaping immune control (Figure 2D-E). These associations would suggest KLRG1 ^+^ Tconv possess a tumor-supportive role in the treatment-naïve setting. To determine whether KLRG1 ^+^ Tconv also associate with response to ICB, we compared frequencies of KLRG1 ^+^ Tconv in MC38-bearing mice undergoing progressive growth versus those undergoing curative responses to anti-PD-1 (Figure 2F). While KLRG1 expression was rare on CD4 Tconv on day 7, we observed a significant decrease in KLRG1 ^+^ Tconv cells in anti-PD-1 treated animals on both day 10 (p=0.014) and day 13 (p=0.0033; Figure 2H) as compared to isotype treated animals. In total, these data indicate KLRG1 ^+^ Tconv frequencies associate consistently with tumor burden over multiple time points throughout tumor progression and across distinct models, and may have unappreciated functional roles in the TME or utility as biomarkers of response to immunotherapy.

**Figure 2.**
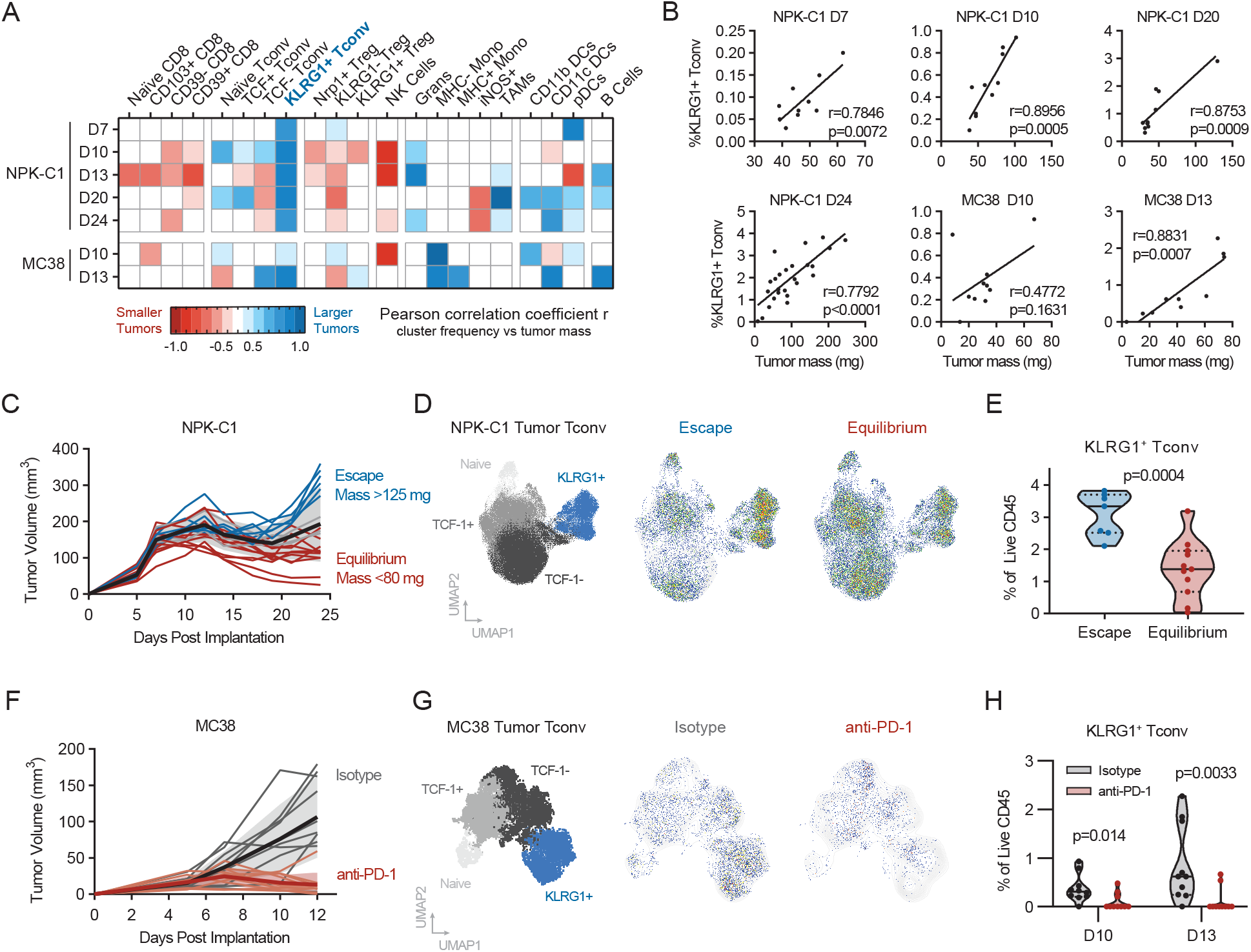
Association between KLRG1+ Tconv frequency and tumor progression. (A) Heatmap of Pearson correlation coefficients comparing metacluster frequency (% of CD45) versus tumor mass. (B) Dot plots of selection correlations as shown in (A). (C) Schematic of NPK-C1 model with colors representing escaping tumors (>125mg) versus those remaining in equilibrium (<80mg) on D24. (D) UMAP projections of CD4 Tconv metaclusters and pseudocolor representations of cells from Escape tumors versus Equilibrium tumors. (E) Violin plot of KLRG1+ Tconv frequency in Escape versus Equilibrium tumors. (F) Schematic of MC38 model. (G) UMAP projections of CD4 Tconv metaclusters and pseudocolor representations of cells from isotype versus anti-PD-1 treated tumors. (H) Violin plot of KLRG1 + Tconv frequency in isotype versus anti-PD-1 tumors on D10 and D13. Data were generated from one experiment of n=10-25 mice per/condition and time point.

### Tumor Helios^-^KLRG1^+^ Treg deficiency associates with enhanced tumor control

Based on these findings we also investigated whether KLRG1 ^+^ Treg phenotypes associate with tumor progression or ICB response. While KLRG1 ^+^ Tregs as a bulk population did not correlate consistently with tumor burden in the NPK-C1 or MC38 models (Figure 2A), we observed two Treg sub-clusters defined by Helios^-^KLRG1^+^ that correlated significantly with tumor burden on day 24 in the NPK-C1 model, which indicates an association between this Treg phenotype and transition of NPK-C1 tumors from equilibrium to escape (Figure 3A-E). To test the generalizability of this Treg phenotype and its functional associations across models, we asked whether the Helios^-^ KLRG1^+^ Treg phenotype is associated with curative responses to immunotherapy in MC38 (Figure 3F). FlowSOM clustering produced two Helios^-^KLRG1 ^+^ Treg clusters that were present in progressive untreated tumors, but were significantly lost on day 10 (p=001) and day 13 (p=0.00017; Figure 3H) in anti-PD-1 treated animals. Therefore, the Helios^-^ subset of KLRG1 ^+^ Tregs exhibits relevant associations with disease progression and response to ICB in distinct tumor models, suggesting a potentially generalizable role for Helios^-^KLRG1 ^+^ Tregs in tumor immunity and response to immunotherapy.

**Figure 3.**
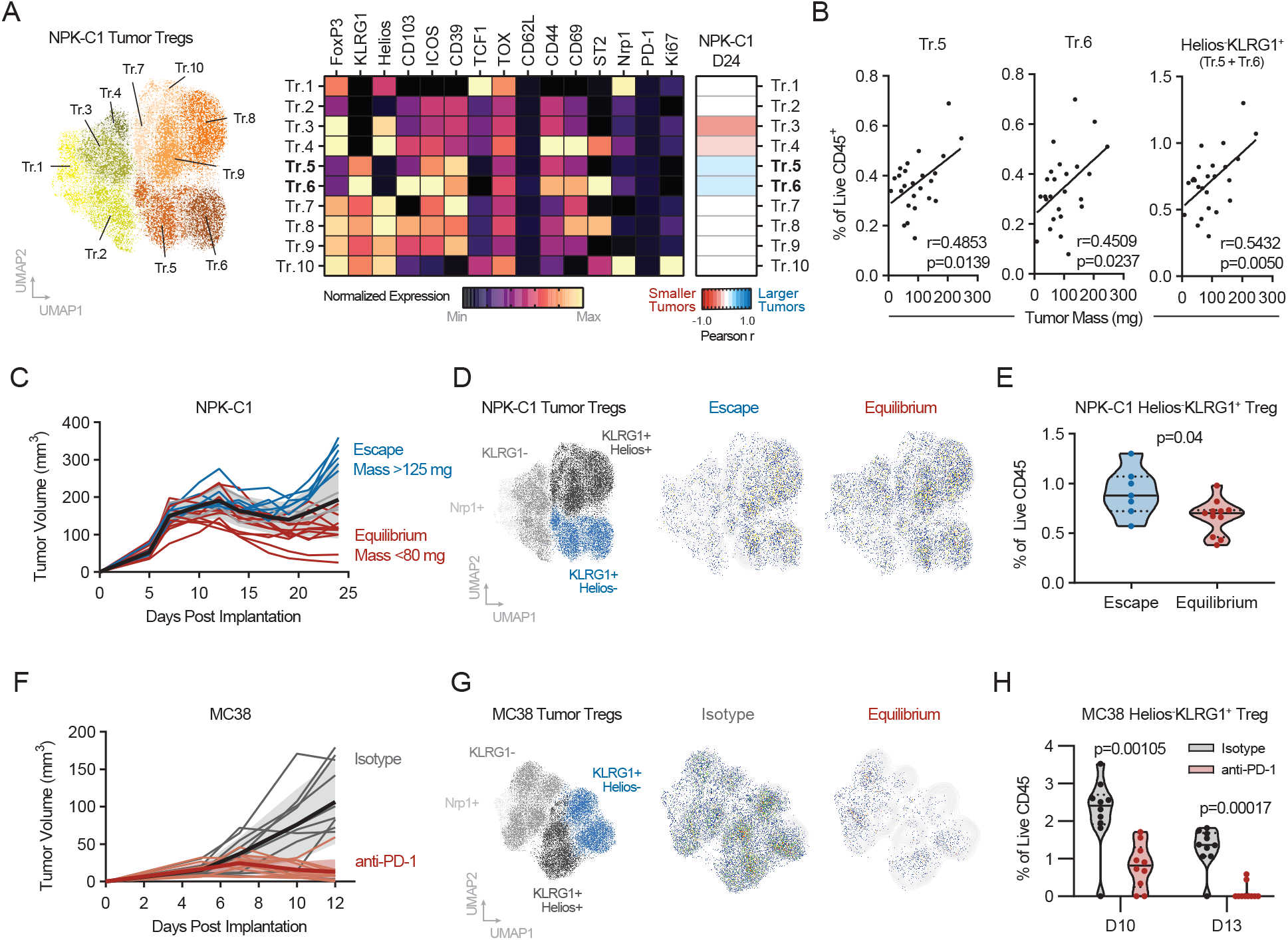
Association between Helios-KLRG1+ Treg frequency and tumor progression. (A) UMAP projection and FlowSOM clustering of NPK-C1 tumor Tregs, and cluster heatmap of normalized phenotypic marker expression. Pearson correlation of each cluster frequency versus tumor mass shown by heat map, and selected correlation dot plots in (B). (C) Schematic of NPK-C1 model with colors representing escaping tumors (>125mg) versus those remaining in equilibrium (<80mg) on D24. (D) UMAP projections of CD4 Treg metaclusters and pseudocolor representations of cells from Escape tumors versus Equilibrium tumors. (E) Violin plot of Helios-KLRG1 + Treg frequency in Escape versus Equilibrium tumors. (F) Schematic of MC38 model. (G) UMAP projections of CD4 Treg metaclusters and pseudocolor representations of cells from isotype versus anti-PD-1 treated tumors. (H) Violin plot of Helios-KLRG1+ Treg frequency in isotype versus anti-PD-1 tumors on D10 and D13. Data were generated from one experiment of n=10-25 mice per/condition and time point.

### KLRG1+ CD4 T cells correlate with tumor progression in kidney cancer

To validate the translational relevance of our findings, we asked whether KLRG1 in human tumor-infiltrating T cells associates with clinical disease progression. For this, we analyzed data from two cohorts of clear cell renal carcinoma (ccRCC) patients whose tumors were profiled by single cell RNA sequencing (10,11). At the gene expression level, *Klrg1* is present largely in CD8 T cells as well as CD4 and NK cells (Figure 4A).

**Figure 4.**
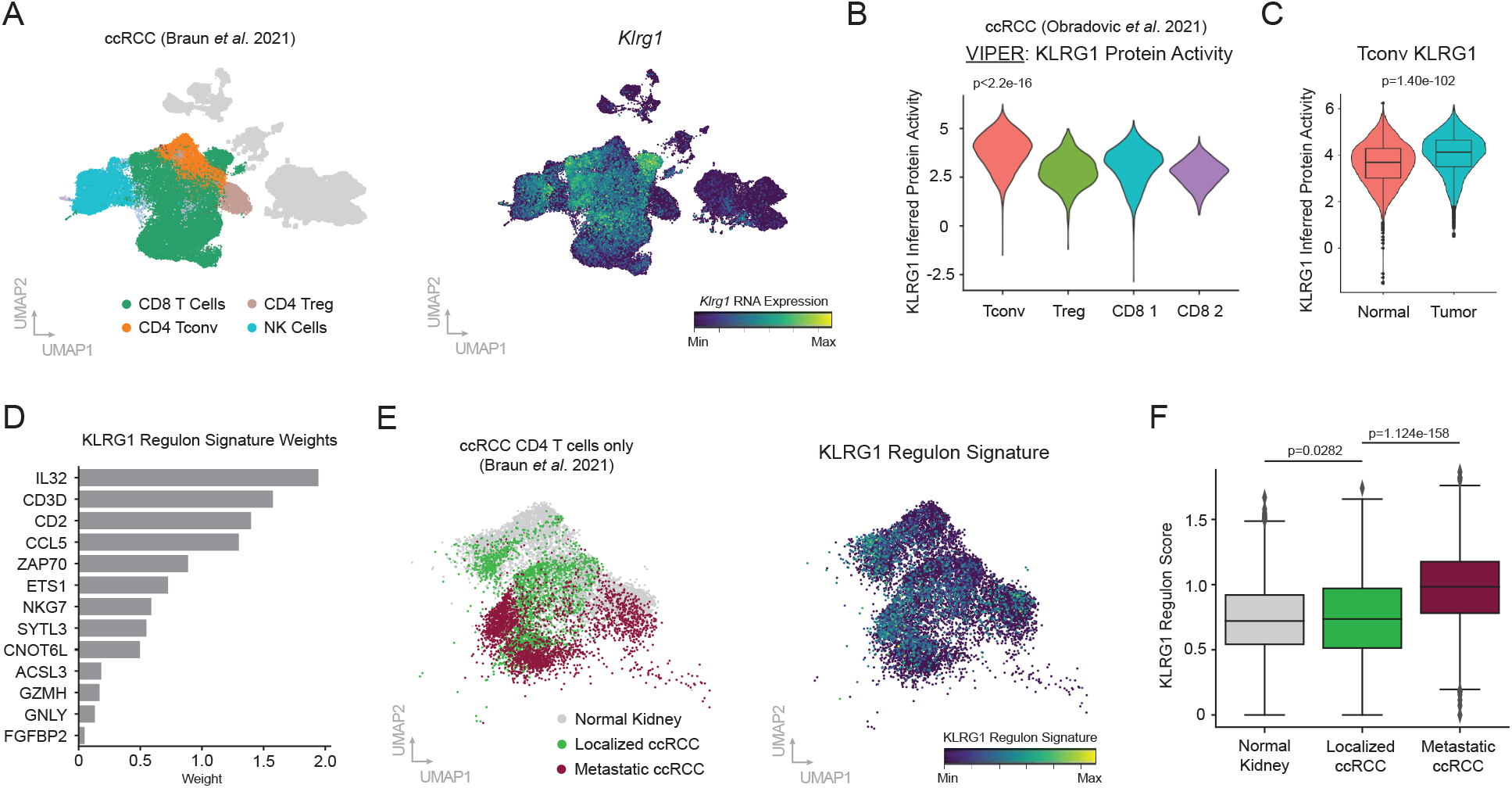
Validation of KLRG1+ Tconv association with tumor progression in human RCC. (A) UMAP projection of ccRCC single cell data from Braun, *et al*. 2021. *Klrg1* gene expression is shown. (B) VIPER inferred protein activity of KLRG1 in ccRCC T cell populations from Obradovic, *et al*. 2021. (C) Inferred activity of KLRG1 in CD4 Tconv in ccRCC tumors versus adjacent normal kidney. (D) Bar plot of gene weights learned by random forest regression for KLRG1 regulon signature score. (E) UMAP projection of ccRCC CD4 T cells from Braun, *et al*. 2021. KLRG1 signature score is shown. (F) Quantification of KLRG1 regulon score in CD4 T cells in normal kidney versus localized ccRCC versus metastatic ccRCC.

However, we inferred KLRG1 protein activity by VIPER (Virtual Inference of Protein activity by Enriched Regulon analysis; see (11,13) for validation of method) and found by contrast that KLRG1 inferred protein activity is highest in CD4 Tconv in RCC tumors (Figure 4B). KLRG1 protein activity is significantly increased in Tconv infiltrating ccRCC as compared to Tconv in adjacent normal kidney parenchema, further validating the relevance of the KLRG1+ CD4 T cell phenotype in human cancer (p=1.0e-102; Figure 4C). To test whether KLRG1 activity in CD4 Tconv associates with disease progression, we computed a KLRG1 regulon signature consisting of genes whose upregulation at the RNA level predicts KLRG1 protein activity (Figure 4D), then asked whether this KLRG1 signature is enriched in CD4 Tconv infiltrating late-stage vs early-stage ccRCC. For this, we focused on the Braun dataset as it was generated to profile T cell phenotypes across stages of ccRCC progression (10). We found the KLRG1 regulon signature score increased from naïve kidney to localized ccRCC (p=0.0282), and was highly enriched in metastatic disease (p=1.124e-158; Figure 4E-F). These data validate our preclinical flow cytometry observations and warrant additional mechanistic and validation studies to understand the utility of tumor-infiltrating KLRG1^+^ CD4 T cell phenotypes as biomarkers and/or targets of cancer immunotherapy.

## Discussion

In this study, we generated a comprehensive spectral flow cytometry dataset of productive versus non-productive immunity across tissues and relevant time points in the NPK-C1 and MC38 murine tumor models. Using dimensionality reduction and minimally-supervised clustering analyses, we identified sub-phenotypes of tumor-infiltrating CD4 Tregs and Tconv cells marked by KLRG1 expression, whose relative frequencies associate with tumor burden, tumor progression from equilibrium to the escape, and response to anti-PD-1 immunotherapy.

CD4 T cells expressing KLRG1 have been previously discovered in both murine and human tumors, yet the etiology and functional properties of these cells remain incompletely described(14). In a related study by Li, *et al*. KLRG1^+^Amphiregulin^+^ Tregs were shown by single cell RNA sequencing and flow cytometry to accumulate in autochthonous Kras^G12D^ p53^-/-^ (KP)-driven lung tumors. These Tregs were driven by IL-33 signaling, thus Treg-conditional ablation of ST2/IL-33R reduced KLRG1 ^+^ Treg frequency and de-repressed CD8 T cell mediated antitumor immunity(12). While our data do not disqualify a role for IL-33 in inducing KLRG1 ^+^ Tregs or Tconv cells in our models, we note that ST2/IL-33R expression was not characteristic of or restricted to KLRG1^+^ CD4 T cells in our data. It remains possible other tissue factors such as TGF-β contribute to the KLRG1^+^ Treg or Tconv in the TME, as observed in the intestinal environment(15). Our data supports previous observations that KLRG1 ^+^ Tregs and Tconv exhibit an activated, terminally differentiated phenotype expressing high levels of CD39 and ICOS(16,17). While our correlative data suggest these phenotypes are tumor-supportive, additional studies are warranted to determine the mechanisms by which this occurs. For KLRG1 ^+^ Tconv, it is possible that these cells are exhausted, analogous to a KLR^+^ exhausted CD8 T cell program described in the chronic LCMV model(18). Alternatively, KLRG1+ Tconv may take on a suppressive function as has been noted *in vitro(19*), or as a result of high CD39 expression as seen with exhausted CD8 T cells in cancer(20). However, it remains possible KLRG1 ^+^ Tregs are a passive byproduct and quantifiable marker of an alternative dominant suppressive factor in the TME. Additionally, while KLRG1 ^+^ Tregs are known to be highly immunosuppressive(16,21), our data suggest it is a Helios^-^ subset of KLRG1 ^+^ Tregs that may be uniquely functionally relevant, as they are a highly consistent marker of tumor burden, progression, and ICB response in our models. Whether these Tregs are peripherally induced from KLRG1^+^ Tconv, and if these cells exhibit unique functional characteristics from Helios^+^KLRG1^+^ Tregs are open questions for further study.

In total, this study calls for future investigation into KLRG1^+^ CD4 T cell subsets as potential targets of – or putative biomarkers for – cancer immunotherapy. Though our data warrant mechanistic follow-up and additional validation in clinical specimens, the concordance of our observations with others in various other ectopic and autochthonous models suggests these cell types may be generalizable across tumor types in both mice and humans(12,21). Furthermore, our findings are proof of concept that a longitudinal high-parameter spectral flow cytometry approach can be applied to additional model systems and time points to extract novel targets and/or biomarkers from dynamic “temporal atlases” of antitumor immunity. It is likely these and related approaches will be critical in better understanding the complex, dynamic interactions that shape the TME, and ultimately lead to more effective utilization of immunotherapy in the clinic.

---

**Supplementary Figure 1.**
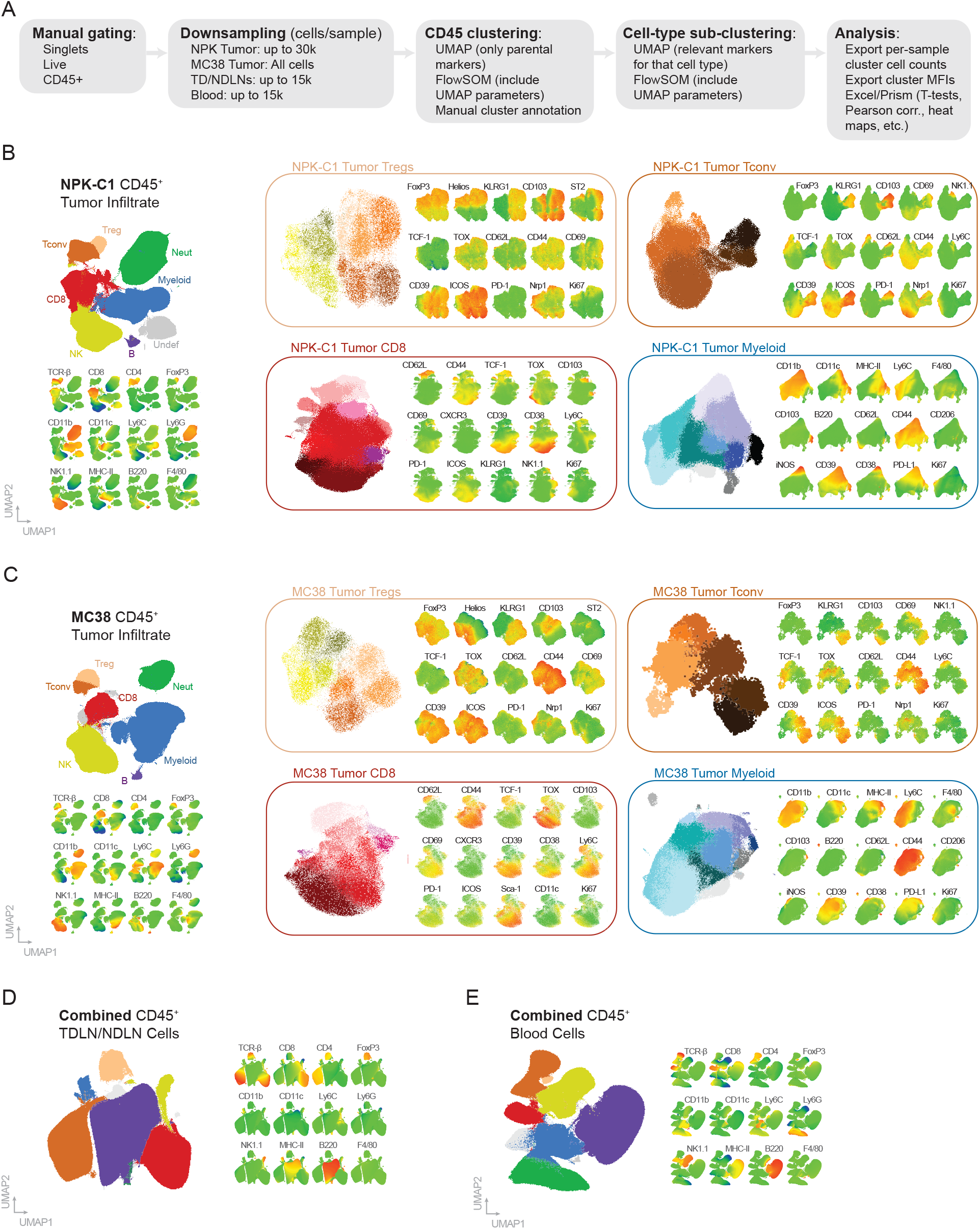
High dimensional analysis workflow and detailed cluster phenotyping. (A) Schematic of flow cytometry analysis workflow. (B) CD45+ clustering, cell-type sub-clustering, and associated phenotypes as represented by heatmap coloring of NPK-C1 infiltrating immune cells. (C) CD45+ clustering, cell-type subclustering, and associated phenotypes as represented by heatmap coloring of MC38 infiltrating immune cells. (D) CD45+ clustering and associated phenotypes as represented by heatmap coloring of all TDLN and NDLN immune cells across both tumor models. (E) CD45+ clustering and associated phenotypes as represented by heatmap coloring of all blood immune cells across both tumor models.

## Acknowledgments

We thank Dr. Cory Abate-Shen at Columbia University for kindly sharing the NPK-C1 cell line.

